## Supplementary material for "Reassessing the modularity of gene co-expression networks using the Stochastic Block Model": SI Figure 3

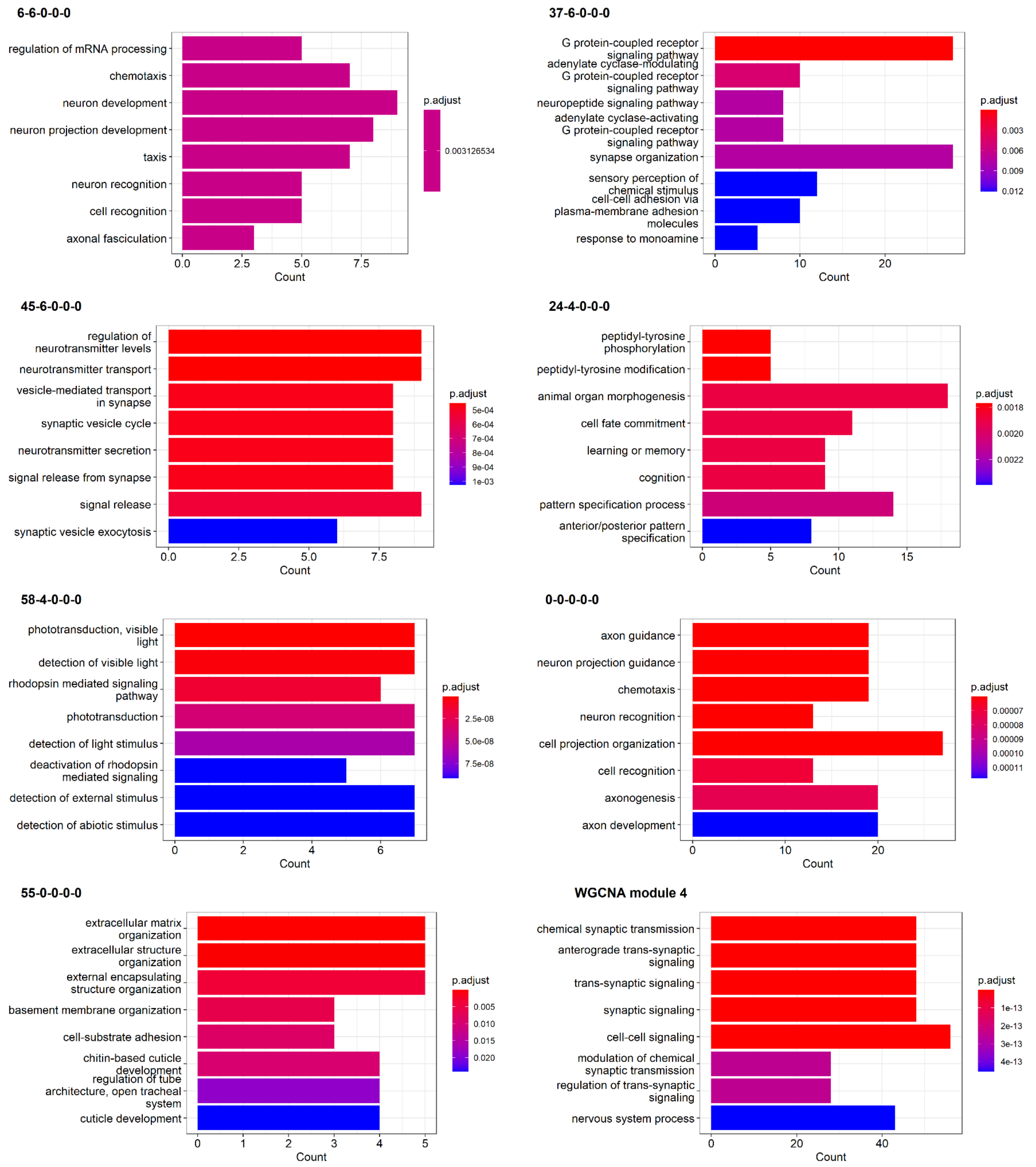


**SI Figure 3**: Enriched GO categories in a level-3 block in the head (0-0-0), related to neural signaling. Panels show the corresponding level-1 blocks. Bars correspond to the top 8 GO categories, the x-axis shows the number of genes associated with each term. The last panel shows the most similar WGCNA module, which also contains signaling-related genes, but at a lower resolution and fails to cluster the phototransduction genes, which are in WGCNA module 5 (not shown, but see SI table 2.1).
