## Supplementary material for "Reassessing the modularity of gene co-expression networks using the Stochastic Block Model": SI Figure 2

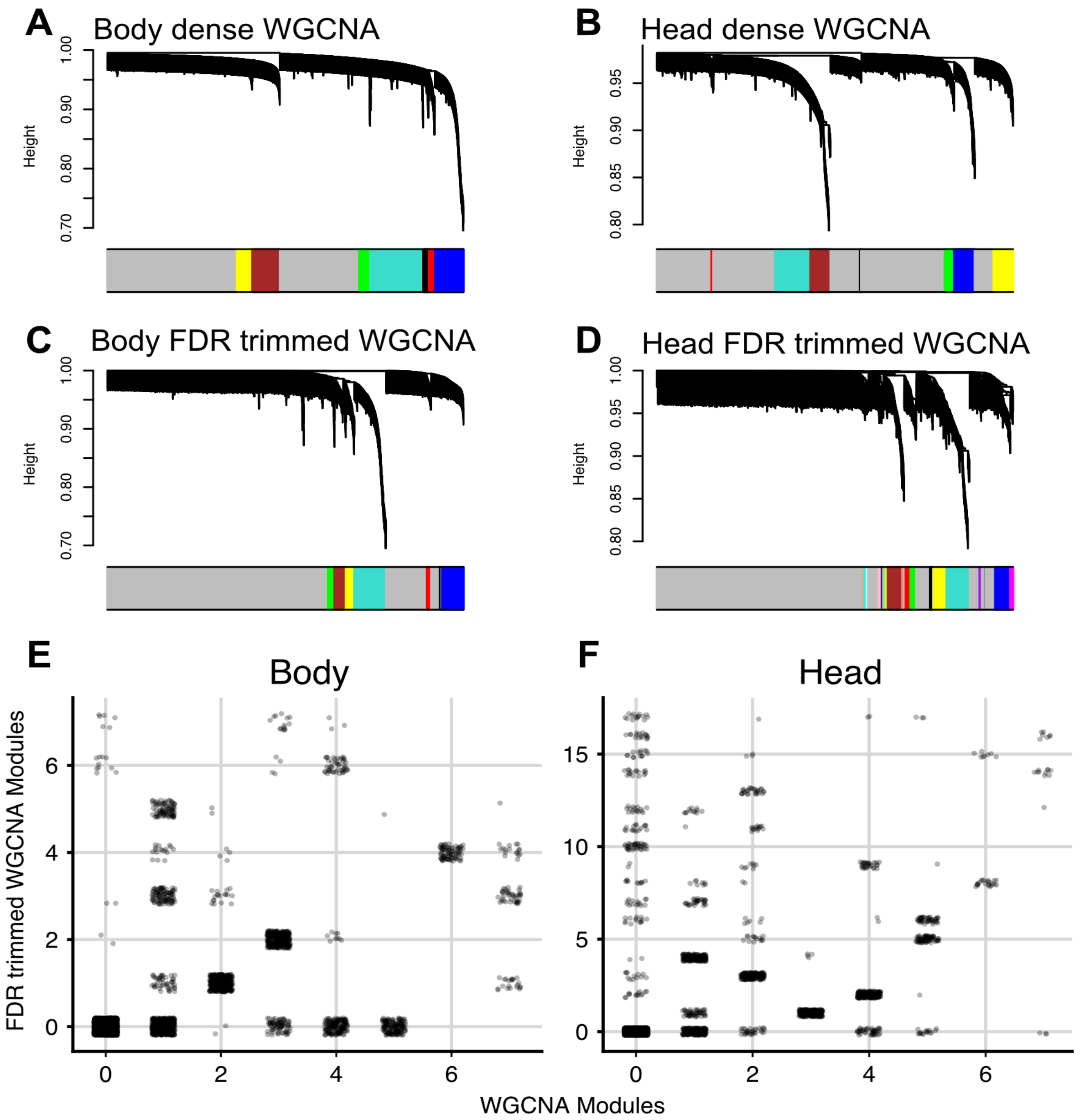


**SI Figure 2**: Comparison of the WGCNA clustering using the standard WGCNA workflow and using the same FDR trimming of non-significant correlations across genes that we use in the SBM clustering. (A) WGCNA clustering of body tissue using the standard workflow. (B) WGCNA clustering of head tissue using the standard workflow. (C) WGCNA clustering of body tissue using FDR trimming of non-significant correlations. (D) WGCNA clustering of head tissue using FDR trimming of non-significant correlations. (E) Correspondence of genes in trimmed and non-trimmed WGCNA, showing the mapping of modules between the two methods. The WGCNA clustering changes slightly when FDR trimming is applied, but a similar number of genes are clustered in both methods. The resulting modules are different but can be reasonably mapped to each other, suggesting that the overall structure of the gene network is largely maintained. This comparison shows that while the specific processing of gene expression correlations can impact the final clustering results, the WGCNA method is relatively robust to these changes and produces comparable modules even when the network is trimmed using FDR correction.
