## Supplementary material for "Reassessing the modularity of gene co-expression networks using the Stochastic Block Model": SI Figure 1

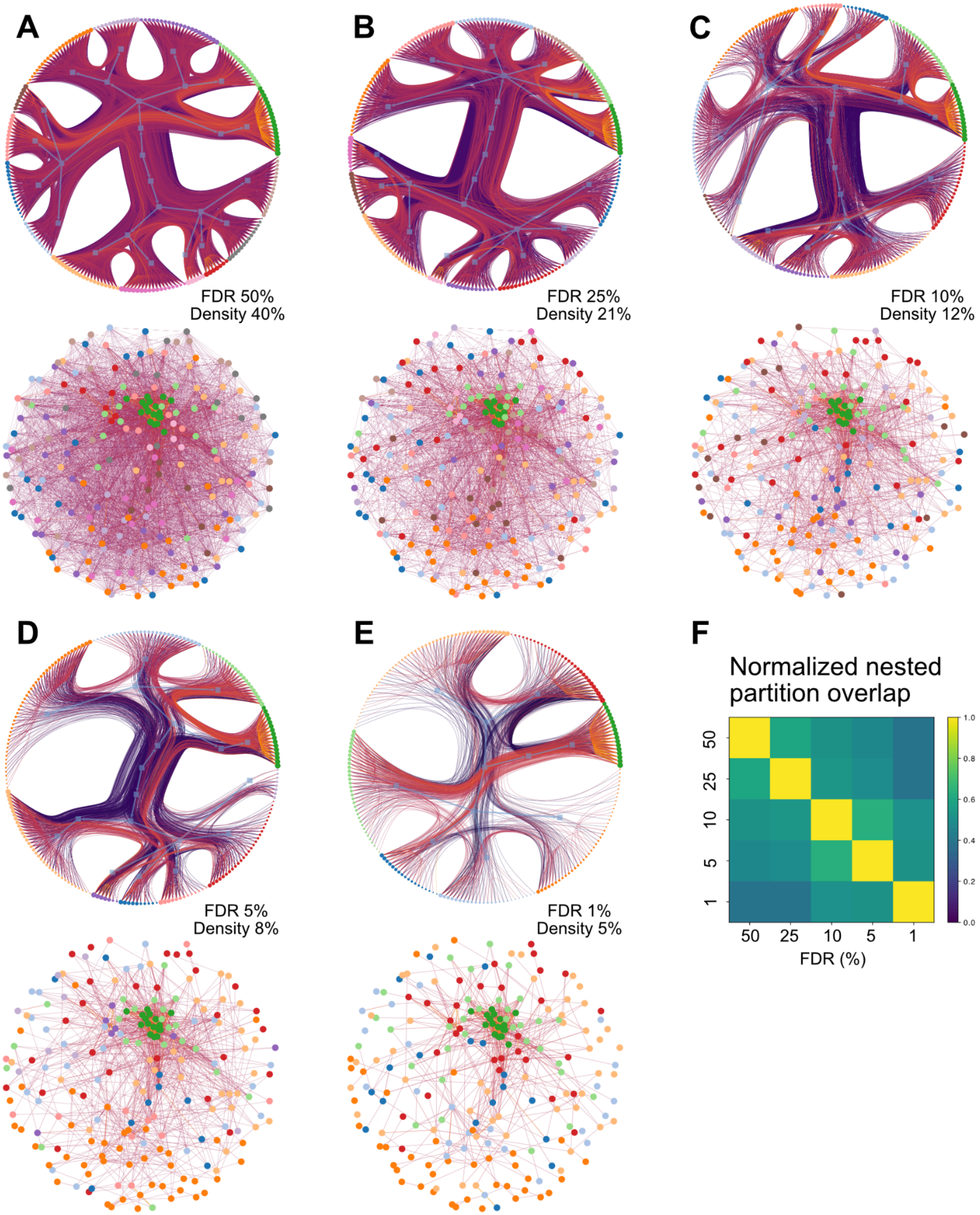

A
FDR 50%
Density 40%
B
FDR 25%
Density 21%
C
FDR 10%
Density 12%
D
FDR 5%
Density 8%
E
FDR 1%
Density 5%
F
Normalized nested partition overlap
50
25
10
5
1
FDR (%)
1.0
0.8
0.6
0.4
0.2
0.0

**SI Figure 1**: Comparison of gene co-expression networks and their corresponding SBM-inferred clusters at different FDR thresholds for a random subset of 300 genes from the head dataset. We trim the networks at different FDR levels (50%, 25%, 10%, 5%, and 1%;A to E respectively ). To compare the networks at all FDR levels directly, we only use the genes that are connected at all FDR levels, resulting in networks with 213 genes. Because this is a random sample of the transcriptome, we do not expect every gene to be meaningfully assigned to a block, and we do not expect the relations between blocks to be completely stable either between different posterior samples of the nested partition. We do, however, expect that true biological blocks be consistent, and that the general structure of the partition remains stable across FDR networks and realizations. For panels A-F, we see the SBM hierarchical clustering (top) and the gene network (bottom). Genes are colored according to their level-1 blocks, and partitions at different FDR values have been optimally aligned to have the same color (Peixoto (2021) 10.1103/PhysRevX.11.021003). Because there are different numbers of blocks at each FDR level, the colors are not always present at every FDR level.

(A) Gene co-expression network at 50% FDR threshold. The network is highly connected and presumably noisier. Even so, the SBM is able to identify a structure that is similar to the other networks. The most apparent difference is that the increase in the number of edges allows for more subdivision of the blocks. The dark green block is clearly visible at this FDR level, but other blocks that are present in the other FDR values appear more subdivided here. It's difficult to determine whether this subdivision is warranted, or if it is a consequence of the lower signal-to-noise ratio in this more highly connected network.

(B) Gene co-expression network at 25% FDR threshold. The network is less connected than the 50% FDR network, and the blocks that are identifiable at lower FDR values become more apparent. In particular, the green assortative block remains as the most prominent, with the light green (around the green block) and orange (lower part of all the networks) blocks also becoming discernible.

(C-E) Gene co-expression networks and their corresponding SBM-inferred clusters at 10%, 5%, and 1% FDR thresholds, respectively, for the same subset of genes. The network connectivity decreases as the FDR threshold becomes more stringent, but several blocks remain stable and the structure of the hierarchy remains similar.

(F) A matrix of nested partition overlap across all FDR values calculated using the graph_tool function *nested_partition_overlap*. These values are scaled to be between [0, 1] and reflect the degree that the nested partitions match at all levels. See Peixoto (2021) 10.1103/PhysRevX.11.021003 for details. The nested partitions are broadly similar across all FDR values, with most overlap values being above 0.5. Unsurprisingly, networks with more similar FDR values show more similar partitions.

The stability of the inferred blocks across different FDR thresholds demonstrates the ability of the SBM approach to identify robust gene co-expression clusters, even in the presence of varying levels of noise and connectivity in the network. This highlights the potential of the SBM method for uncovering biologically relevant patterns in gene co-expression data, and its resilience to the choice of FDR threshold used for network construction, provided that the threshold is sufficiently stringent to remove the more noisy edges.
